# Identification of TAP2 protein variants resistant to inhibition by the HSV1 ICP47 protein

**DOI:** 10.1101/2024.11.18.624061

**Authors:** Gregory Nagy, Rebecca Presley, Shruti Dhar, Hyeongseon Jeon, Lynn Moreira, Olivia Tull, Arkobrato Gupta, Dongjun Chung, Philip N. Tsichlis, Jeffrey D. Parvin

**Author notes:** These authors contributed equally (Gregory Nagy, Rebecca Presley).

## Abstract

Herpes Simplex Virus 1 evades the host immune system by expressing a protein, ICP47, that binds to and inhibits the heterodimeric Transporter Associated with Antigen Processing (TAP). We screened a library of 1786 variants in TAP2, one of the components of the TAP heterodimer, and identified 39 variants that were resistant to inhibition by ICP47. Of these 39 variants, five were individually tested, and three (T257I, S274H, and T244R) were confirmed to be significantly resistant to inhibition by ICP47. This resistance to inhibition did not extend to the Epstein Barr Virus BNLF2a protein, another viral factor known to inhibit antigen presentation by targeting TAP. These three residues localize close to the binding site of ICP47, on the 3D structure of TAP, but only Ser274 is spatially close to the antigenic peptide binding site of TAP. These results functionally resolve the TAP2 residues required for peptide binding from those required for ICP47 binding and identify TAP2 residues whose targeting with small molecule inhibitors could effectively prevent Herpes virus downregulation of antigen processing.

**AUTHOR SUMMARY:** T cells detect virus infected cells via viral peptides in complex with the major histocompatibility complex type I (MHC-I) on the cell surface. Herpes Simplex Virus 1 expression of the ICP47 protein allows the virus to evade detection and clearance of infected cells by host T cells. ICP47 binds to the TAP1/TAP2 protein complex and prevents the loading of antigenic peptides onto the MHC-I protein complex and thereby blocks the antigen processing. In this study, we screened for and identified variants of TAP2 with single amino acid substitutions that are no longer inhibited by ICP47. TAP2 proteins with the identified amino acid substitutions functioned normally in antigen processing even in the presence of ICP47. Results from this study create the potential strategy for drugs that target the identified portions of the TAP2 protein and block ICP47 inhibition of antigen processing while retaining normal TAP1/TAP2 function and thus suggest a new antiviral therapeutic approach for herpes virus infections.

## INTRODUCTION

Clearing Herpes Simplex virus-1 (HSV-1) infected cells depends on the recognition by CD8+ T cells of viral peptides bound to major histocompatibility complex type I (MHC-I) on the surface of infected cells. The CD8+ T cells recognize foreign peptides in complex with host MHC-I, triggering the lysis of the infected cells [1]. One of the key components of the antigen presentation machinery is TAP, the Transporter Associated with Antigen Processing, a heterodimer of two proteins (TAP1 and TAP2). The heterodimeric TAP is an ABC class channel, which transports antigenic peptides from the cytoplasm to the endoplasmic reticulum (ER), where it presents it to MHC-I [2–4]. TAP is required for antigen presentation, as evidenced by the fact that genetic loss of either the TAP1 or the TAP2 subunit results in severe immunodeficiency (Bare Lymphocyte Syndrome) [5,6].

Cellular proteins are processed by the proteasome, generating cytoplasmic peptides eight to thirteen residues in length. These peptides bind the cytoplasmic side of the TAP transporter, which is anchored in the membrane of the endoplasmic reticulum (ER). Following binding, and in the presence of ATP, TAP changes its conformation from inward cytoplasmic-facing to outward ER lumen-facing and releases the peptide into the ER lumen. Following processing by the ER proteins Tapasin, TAPBPR, calreticulin, ERp57, ERAP1, and ERAP2, the peptide is loaded onto MHC-I, and the peptide/MHC-I complex re-localizes to the cell surface, where it is presented to the T cells [1,2,4]. In the absence of TAP function, few antigenic peptides are transported to the ER lumen, and the MHC-I complex remains localized in the ER and does not translocate to the cell surface. Thus, in the absence of TAP function, there is a decrease in MHC-I on the cell surface.

Herpes viruses have developed mechanisms to evade this immune surveillance pathway; Herpes Simplex Virus 1 (HSV1) expresses ICP47, which binds to, and inhibits the function of the TAP peptide transporter [7,8]. The 3D structure of ICP47 in complex with the TAP1-TAP2 heterodimer has been determined. According to this structure, ICP47 binds to the inner channel of TAP and appears to function as a cork blocking peptide transport [9,10]. In the absence of normal antigenic peptide transport, the MHC-I remains intracellular and is not localized to the cell surface, preventing the recognition of HSV1 infected cells by cytolytic CD8+ T cells.

In this study, we screened a multiplexed library of TAP2 variants, to identify those variants that are resistant to inhibition by ICP47. Binding of ICP47 to TAP results in loss of TAP transporter activity. TAP2 variants resistant to ICP47 inhibition were variants that continued to support the expression of MHC-I on the cell surface, even in cells expressing ICP47.

## RESULTS

### Experimental model for the identification of TAP2 variants resistant to HSV1 ICP47

The coding sequence for the 88 amino acid residue ICP47 was cloned in the lentivirus vector, pHAGE2-EF1aL-dsRed-W, downstream of the EF1a promoter, the dsRed coding sequence and the P2A translation re-start signal (Figure 1A). Infection of U2OS cells with this lentivirus at an MOI of < 0.5 resulted in dsRed-positive/ICP47-positive cells and dsRed-negative/ICP47-negative control cells that could be distinguished by flow cytometry. These cells were stained using an anti-human HLA A/B/C (anti-human MHC-I) antibody conjugated to a fluorescent tag. Flow cytometry revealed that the MHC-I surface staining of the uninfected control (dsRed-negative) cells, was consistently higher than the MHC-I surface staining of the ICP47-expressing dsRed-positive cells (Figure 1B). The histograms of the two populations of cells are shown. The infected and the uninfected control cells coexisted in the same samples, eliminating the potential for variability between them because of unintended differences in culture or assay conditions. Using this system, the results were highly reproducible between experiments.

**Figure 1.**
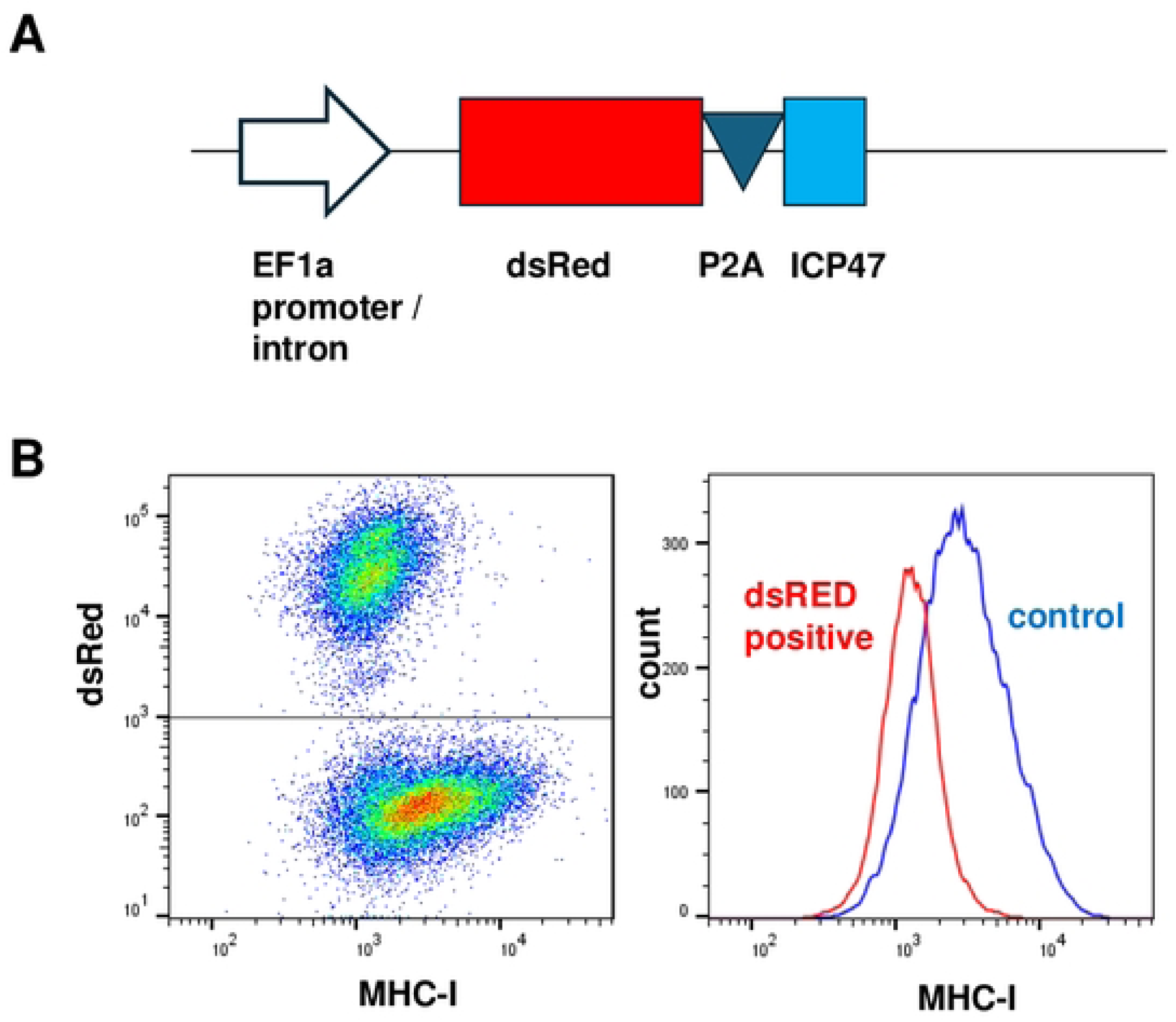
Experimental approach to identifying TAP2 variants resistant to HSV1 ICP47. **A.** The DNA encoding ICP47 was inserted into a lentiviral vector downstream of the EF1a promoter and expressed from a single RNA encoding dsRed, the P2A translation re-start signal, and ICP47. **B.** U2OS cells were infected with the lentivirus described in panel A at an MOI of 0.4. Six days post-infection, cells were stained for MHC-I and analyzed by flow cytometry. The x-axis indicates the intensity of anti-MHC-I antibody stain and the y-axis the intensity of dsRed stain (*left*) or the histogram of the cells with a given MHC-I stain intensity (*right*).

In an earlier study, we used a multiplex functional assay to screen a barcoded TAP2 variant plasmid library for variants defective in the transport of antigenic peptides to the ER (manuscript submitted [11]); this analysis was done in the absence of a viral inhibitor of the process, such as ICP47. The library contained variants in 96 codons (total number of variants 1786) that were expressed in U2OS cell in which the TAP2 gene was knocked out using CRISPR [11]. These 96 codons included residues known to be functionally important for TAP, residues known to be subjected to post-translational modification (PTM), and conserved serines, threonines, and tyrosines that are potential acceptors of PTMs. The library of TAP2 variants was in the pUC-19 plasmid backbone that lacked a eukaryotic promoter, but the plasmid contained an attB site upstream of the hygromycin resistance gene, P2A, and the TAP2 library. Transfection of the library of TAP2 variants into U2OS-TAP2-KO cells that harbored a single attP site downstream of the RSV-LTR and the Bsb1 integrase [12], resulted in the integration of a single TAP2 variant per cell via attP/attB recombination. The attP/attB recombination placed the hygromycin resistance-P2A-TAP2 cassette downstream of the tetracycline-regulated RSV-LTR. The analysis of the data in this prior study identified functionally important TAP2 residues, including residues of confirmed or potential phosphorylation sites, whose substitution gave rise to inactive or hyperactive TAP2 variants [11].

In the present study, the U2OS-TAP2-KO cells, transduced with the library of TAP2 variants, were infected with the dsRed-ICP47 expressing lentivirus at an MOI of 0.6, and 5 days later, they were stained with the MHC-I-specific antibody, and analyzed by flow cytometry. The flow cytometry analysis showed that the dsRed-negative/ICP47-negative cells were a mixture of MHC-I positive and negative cells, as expected. On the other hand, the dsRed-positive/ICP47 positive cells, were almost universally MHC-I negative (Figure 2A).

**Figure 2.**
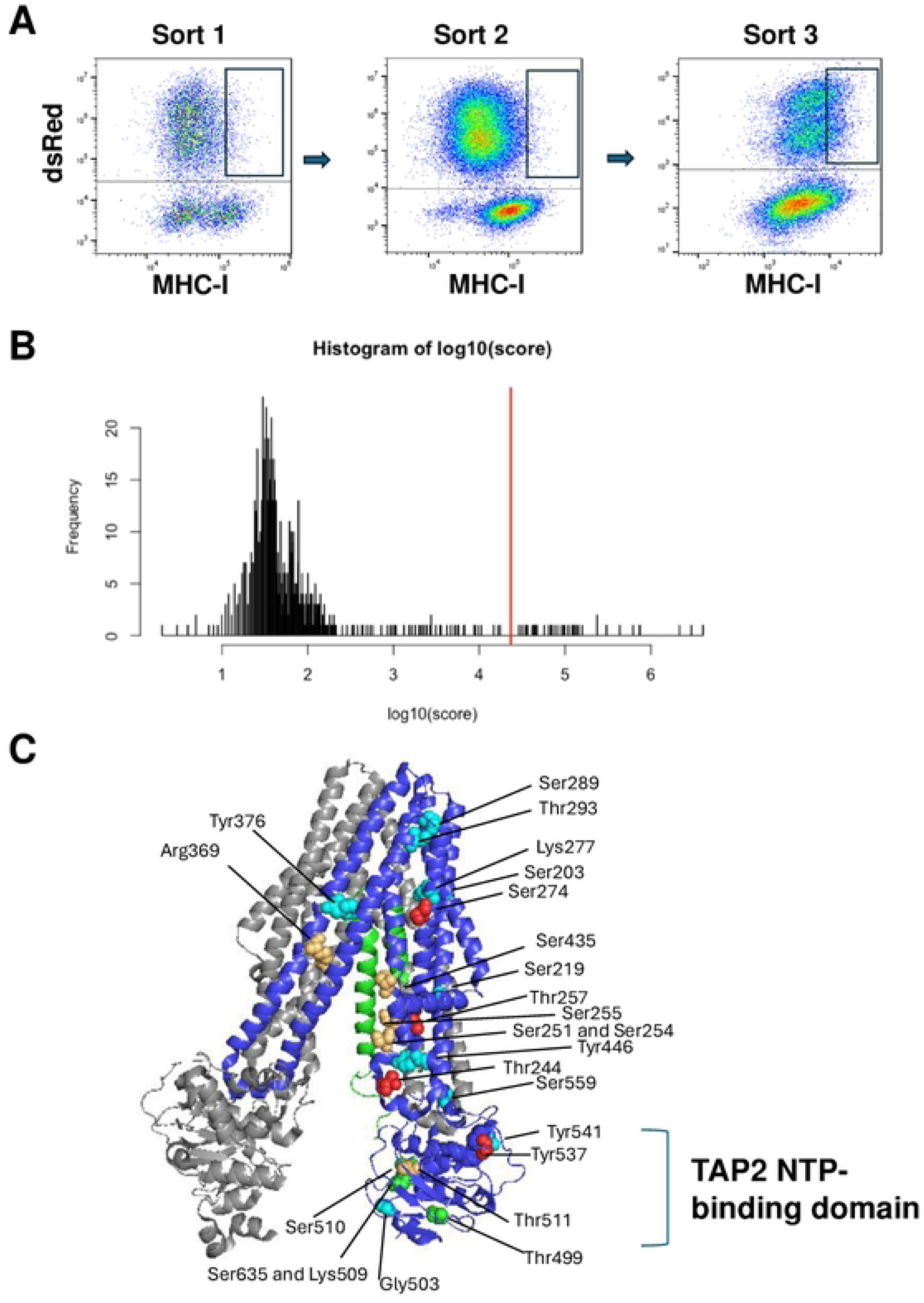
Isolation of TAP2 variants resistant to inhibition by ICP47. **A.** A multiplexed library of variants of TAP2 were infected with the lentivirus expressing dsRed and ICP47 at an MOI of 0.5 and stained with a fluorescent antibody specific for MHC-I (*left*). Cells were collected in the indicated box and re-passaged to amplify the cell number for a second sort. To retain an internal control for Sort 2, uninfected U2OS TAP2-KO cells rescued with wild-type TAP2 were included as 30% of the sample (*middle*). DsRed-positive cells with high MHC-I were collected, indicated by the box, and re-passaged to amplify the cell number for the third sort. For Sort 3, cells marked with high abundance of MHC-I were collected, indicated by the box (*right*), and genomic DNA containing the barcode sequence was amplified and sequenced. **B.** Histogram of the frequency of different TAP2 variants detected in the cells from Sort 3 with high expression of MHC-I surface stain. The red line indicates the divider between variants detected a significantly high frequency to the right and variants that were not significantly enriched in the sample (to the left). **C.** The 3D structure of TAP1 (gray), TAP2 (blue), and ICP47 (green) is shown. Residues of TAP2 at which variants were resistant to inhibition by ICP47 are indicated with the most frequently observed residues in red, next most frequently observed residues in orange-yellow, and lesser observed residues in green or blue. The TAP2 NTP-binding domain is indicated, and the similar domain in TAP1 is not labeled but is directly across from this domain on the left.

We reasoned that the low number of ICP47-expressing cells with high abundance of MHC-I on the cell surface represented two classes of cells: 1) those expressing TAP2 sensitive to ICP47 which stochastically express MHC-I at levels that place them at the upper end of the population of cells expressing low MHC-I; or 2) those expressing a TAP2 that was resistant to ICP47. Upon reanalysis, the cells in the first group would be expected to shift to the left, while the cells in the second group would be expected to continue expressing high MHC-I on the cell surface. Repeated sorting of the MHC-I-positive cells by flow cytometry and selection of cells with high levels of MHC-I on the surface would eventually enrich for the cells expressing variants of TAP2 resistant to ICP47. To test this hypothesis, we collected the cells from the library with high abundance of MHC-I surface expression (see box in Figure 2A, *left*). After expansion of these cells, they were mixed with U2OS-TAP2-KO cells rescued with wild-type TAP2; these latter cells did not express the dsRed-ICP47 proteins and were readily detected as a separate population on flow cytometry. 70% of the cells were from the library of TAP2 variants and 30% of the cells were from WT-TAP2 expressing cells. The 70/30 mixture of cells was stained for MHC-I surface expression. On the second round of flow cytometry analysis, there was no evident enrichment for cells with high expression of MHC-I on the cell surface, but the cells in that portion of the spectrum were gated and collected. On the third repetition of this analysis, the bulk of the dsRed-positive cells were shifted to higher levels of MHC-I expression, indicating that we had successfully selected a cell population transduced with TAP2 variants that were resistant to ICP47. We collected the high MHC-I surface expressing cells from this third sort, and we sequenced from the genomic DNA the barcode integrated in the cell genomes that identified the TAP2 variant in each cell.

We recovered 811 TAP2 variants from this pool of cells expressing high levels of MHC-I and therefore resistant to ICP47 (Table S1). The histogram (Figure 2B) shows the distribution of read-counts for each TAP2 variant in the cells with high expression of cell surface MHC-I. We utilized a conservative statistical approach (see Methods) to model the occurrence of variants in the pool of cells, and the 39 variants with the highest frequency in the pool were significantly enriched (P-value=0). The remaining 772 variants were not significantly enriched in the population (P-value=1). The average number of reads for the 39 high MHC-I expressing variants was 381,299 reads per variant with a median value of 108,454 reads per variant. By contrast, the remaining 772 variants had an average number of read counts of 283 reads per variant and a median of 44.5 reads per variant (Table S1).

Our prior studies [11] addressed the regulation of MHC-I cell surface abundance by TAP2 variants, independent of ICP47. Here, we applied these data to examine the ICP47-independent functional classification of the 39 ICP47-resistant variants (Table S2). Of the 39 ICP47-resistant variants, 33 were classified as functional, five were classified as loss of function (LOF) (Y537P, S510F, K509Q, G503V, Y541G) and one was classified as hyperfunctional (T293P). We were of course surprised to observe that some of the ICP47-resistant variants had been classified as LOF in our original screen. One possible explanation was that the cells expressing these variants were disproportionately abundant in the original cell pool, prior to selection, and that they were carried during the selection by binding to cells transduced with truly ICP47-resistant variants. To test this hypothesis, we compared the frequency of each variant among the total pool of variants and ranked the missense variants from 1 to 1252 based on the read count frequency of each variant (Table S2) [11]. The top-ranked LOF variant, Y537P, was the 24^th^ most abundant variant in the pool, and perhaps that did contribute to it being the sixth-most abundant ICP47-resistant variant. On the other hand, the other LOF variants that scored as ICP47-resistant were not especially abundant with ranks ranging from 244 (G503V) to 975 (S510F). Thus, the presence of LOF variants among the ICP47-resistant variants could not be attributed to anomalously high representation in the pool of variants. Two other possibilities for the appearance of LOF variant among the ICP47-resistant variants were that these variants were misclassified in the original screen or that the interaction with ICP47 caused the conversion of LOF into functional. Results described below with the singleton analysis of selected variants excluded these two possibilities for the presence of LOF variants among the ICP47-resistant variants.

### Structural considerations of TAP2 ICP47-resistant variants

The structure of the TAP1/TAP2 heterodimer bound to ICP47 has been determined by cryo-electron microscopy [9,10]. In Figure 2C, TAP1 was shown in gray, TAP2 residues 130-681 in blue, and ICP47 residues 1-55 in green. The 39 ICP47-resistant TAP2 variants were generated by substitutions in 26 residues, of which 24 were mapped on the 3D structure of the complex. The six most abundant variants in the pool were shown on the 3D structure in red, the next six variants in yellow-orange, and the rest in green and blue (Figure 2C). For the most part, the substitutions in the ICP47-resistant variants localized to TAP2 domains that were spatially close to ICP47. For example, Ser274, the residue with the second-highest scoring variant in this experiment (S274H), mapped near one of the two alpha helices of ICP47. Similarly, Ser254 mapped near one of the ICP47 alpha helices, and Thr244 mapped near the ICP47 random coil from residues 46-51. The most commonly detected ICP47 resistant variant had a substitution at Thr257, in the same general region with the substitutions of the other ICP47-resistant TAP2 variants (Figure 2C).

Some of the TAP2 substitutions that gave rise to ICP47 resistant variants mapped at positions relatively distant to ICP47. These include the residue Tyr537, which was a proline in the sixth most frequently observed resistant variant and a valine in the ninth most frequently observed resistant variant. Other residues with ICP47-resistant variants that were displaced from the known ICP47 binding site were Tyr541 and Ser289 and Thr293. It is possible that these residues affect the structural conformation of the TAP complex and thereby affect ICP47 binding to the complex.

A cluster of ICP47 resistant variants harbored substitutions in the TAP2 NTP binding domain; the substituted sites include Thr499, Gly503, Lys509, Ser510, and Ser635. Since the peptide binding-induced conformational change of the TAP1-TAP2 heterodimer is fueled by ATP binding, these substitutions might affect the conformation of the complex, and consequently the binding of ICP47. Alternatively, the 33 carboxy-terminal ICP47 residues (residues 56-88), which were not mapped on the structure but were present in this experiment, could potentially bind to the NTP-binding domain; note that the carboxy-terminus of the ICP47 sequences in the cryo-EM structure mapped adjacent to the TAP2 NTP-binding domain.

### Singleton analysis of selected ICP47-resistant variants

To validate the results of the screen, we selected five variants for functional comparison with wild-type-TAP2 in singleton assays. The selected variants include the most enriched variant T257I, the second most enriched variant S274H, and the fourth most enriched variant T244R. The selection of these variants also considered the fact that different substitutions of the same animo acid residues gave rise to variants with similar phenotypes, albeit with lower scores. Thus, other substitutions in Thr274 gave rise to the 5^th^, 33^rd^ and 37^th^ most enriched ICP47-resistant variants, and other substitutions in Thr244 gave rise to the 16^th^, 24^th^, and 31^st^ most enriched variants. In addition to the three high scoring ICP47-resistant TAP2 variants, we also selected for the singleton assays, a low scoring borderline resistant variant (T499D, 13^th^ most enriched), to evaluate the potential limitations of the multiplex screen. We also tested the Y537P variant since it was the sixth-most enriched variant, as well as another substitution at this residue was ninth-most frequently observed, plus, this residue had previously been scored as a LOF variant [11], making this variant worth testing.

The singleton assays in this study were designed similarly to the singleton assays employed to validate the results of our original screen for the function of TAP2 variants, in the absence of ICP47 (manuscript submitted [11]). Plasmids containing promoterless full length TAP2 were integrated by attP/attB recombination into a single site in TAP2 knockout U2OS cells [11]. The integration placed the wild type TAP2 or the TAP2 variants under the control of the tetracycline-dependent RSV LTR promoter. The integration of all the TAP2 constructs in the same single genomic site minimized variation due to differences in expression between variants. In three separate experiments, cells expressing a single variant TAP2 were infected with the dsRed-ICP47 expressing lentivirus at an MOI of 0.5 and analyzed for MHC-I cell surface expression. In cells not infected with dsRed and ICP47-expressing lentivirus, the wild-type and most of the TAP2 variants had equivalent levels of MHC-I cell surface stain (Figures 3A and S1); the exception was the TAP2-Y537P variant, which was classified as a LOF variant [11] and was shifted to lower MHC-I stain (Figure S1, *left*). Infection of cells with the ICP47-expressing lentivirus shifted the MHC-I stain to lower levels in wild-type cells (Figure S1, *right*). By contrast, the cell surface abundance of MHC-I in TAP2 knockout cells rescued with the TAP2 variants S274H, T257I, and T244R was not affected by the expression of ICP47. The TAP2 knockout cells rescued with the TAP2-T499D variant, and the same cells rescued with wild type TAP2 expressed similar levels of MHC-I on the cell surface (Figures 3A and S1). Additionally, ICP47 reduced the abundance of MHC-I in both, although the reduction was marginally smaller in the cells rescued with the T499D variant. Since the T499D variant was the 13^th^ most commonly detected ICP47-resistant variant, this marginal effect on ICP47 resistance suggested that higher confidence in the interpretation of ICP47 resistance should be considered for the more frequently detected variants. The abundance of MHC-I in cells expressing the LOF variant Y537P in the absence of ICP47 was equivalent to its abundance in ICP47 expressing cells indicating that ICP47 did not alter the function of the TAP2-Y537P variant. Thus, this LOF variant was not converted to functional in the presence of ICP47; it is not clear why any LOF variants were detected among the 39 ICP47 resistant variants in the multiplexed screen.

**Figure 3.**
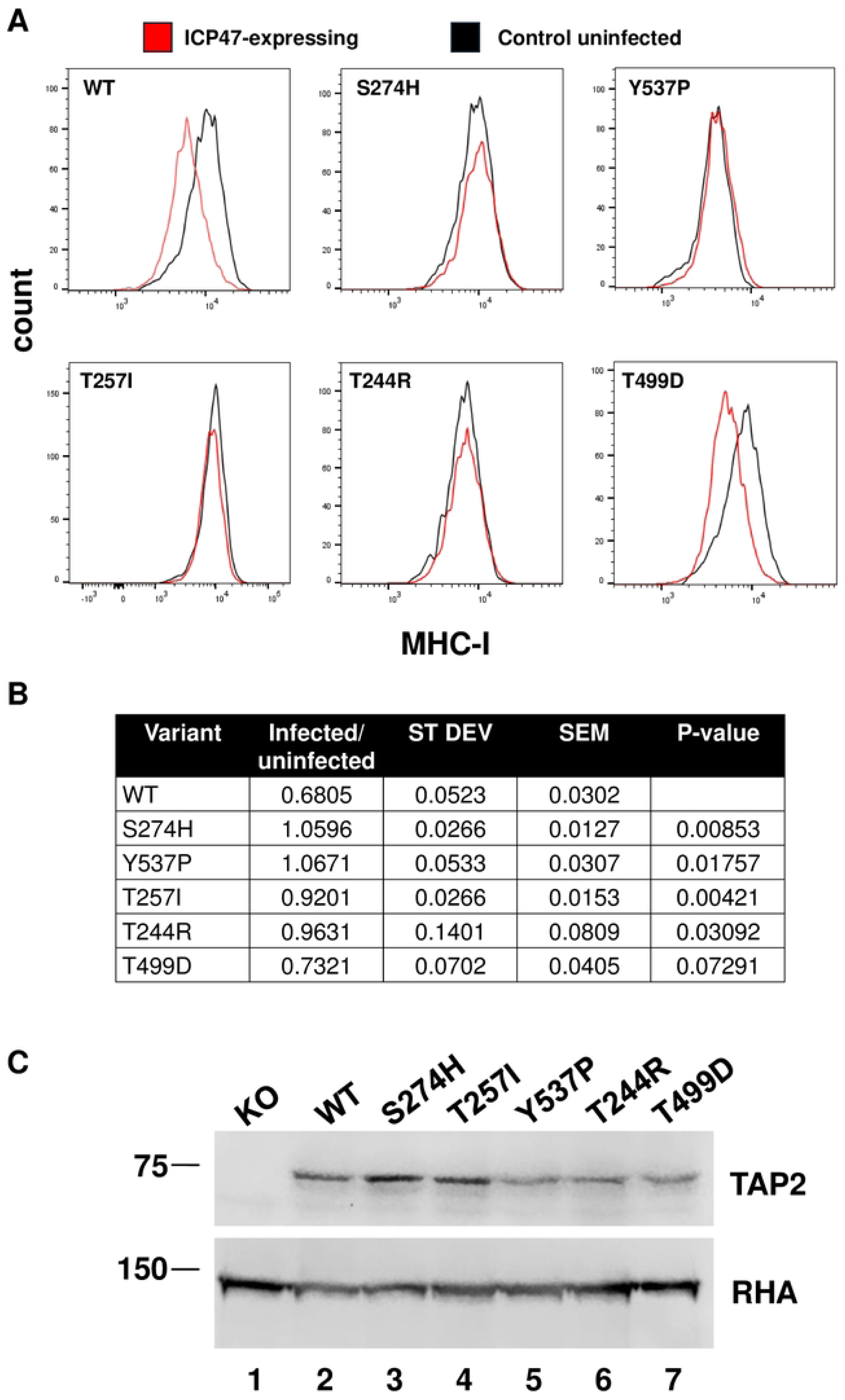
Singleton analysis of TAP2 variants for resistance to inhibition by ICP47. **A.** Histograms indicating the abundance of cells with different intensity of anti-MHC-I cell surface stain in cells expressing ICP47 are indicated for wild-type TAP2, TAP2-S274H, TAP2-Y537P, TAP2-T257I, TAP2-T244R, and TAP2-T499D. **B.** Summary of results comparing the ratio of the median value of the distribution of cells expressing ICP47 divided by the median value of the distribution of control cells. Data were taken from three independent experiments and the average is shown with standard deviation and standard error of the mean. The results of the student’s t-test (two-tailed) comparing each variant to wild-type is indicated. **C.** Immunoblot analysis of TAP2 expression in TAP2-KO cells and the six TAP2 variants analyzed in panel A, as indicated. RNA Helicase A (RHA) was probed as a loading control from the same membrane. Full exposures of the membranes are in Figure S3.

To statistically evaluate the singleton assay data, we measured the median fluorescence intensity (MFI) of cell surface stained MHC-I in the TAP2 knockout U2OS cells rescued with wild type TAP2, or the TAP2 variants, with or without transduction with the dsRed-ICP47 lentiviral constructs. This allowed us to quantitatively measure the ICP47-induced decrease in MHC-I surface expression in cells rescued with wild type TAP2 and each of the TAP2 variants (Table S3). Combining the data from the three biological replicates allowed us to determine the mean value of the ICP47-induced decrease in MHC-I the standard deviation (SD), the standard error of the mean (SEM) and the p value for comparing the variant TAP2-rescued cells to the wild-type TAP2-rescued cells (Figure 3B). The results showed that compared to wild type TAP2, the S274H, T257I, and T244R variants were all significantly resistant to the effects of ICP47 (P < 0.05). The T499D variant was scored as borderline resistant (p= 0.072). The expression of the TAP2 protein in the cells used for the singleton assays was determined by immunoblot analysis, which showed that they were all expressed at levels similar to the wild type TAP2 (Figure 3C).

The Epstein Barr Virus BNLF2a protein is distinct from HSV1 ICP47, but BNLF2a also binds to the TAP complex and inhibits TAP function in antigen presentation [8,13]. To determine whether the ICP47-resistant TAP2 variants were also resistant to BNLF2a, we first cloned the 60 amino acid BNLF2a protein in the pHAGE2-EF1aL-dsRed-W lentiviral vector we had also used for the expression of ICP47. We then infected the TAP2 knockout U2OS cells rescued with wild type TAP2, or with the ICP47-resistant TAP2 variants described above, with the dsRed/BNLF2a-expressing lentivirus, at an MOI of 0.5. The flow cytometry scatter plots (Figure S2) and histograms (Figure 4A) revealed that the wild-type TAP2 and the ICP47-resistant TAP2 variants S274H, T257I, T244R, and T499D were all sensitive to the BNLF2a inhibition of MHC-I surface expression, suggesting that BNLF2a inhibits TAP by binding to different residues than ICP47. In cells expressing both the LOF TAP2-Y537P variant and BNLF2a, the abundance of MHC-I on the cell surface did not further decrease.

**Figure 4.**
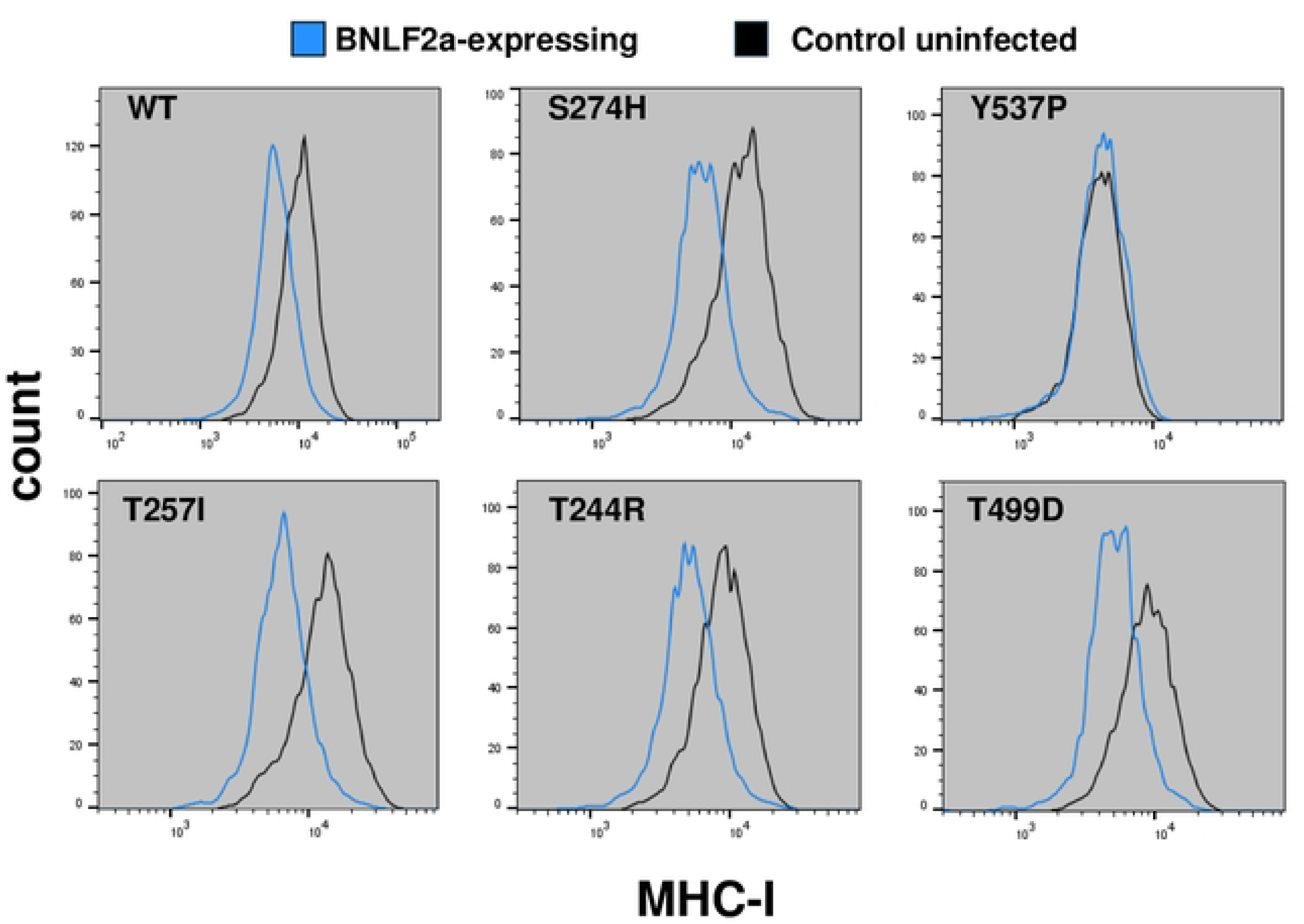
Singleton analysis of TAP2 variants for resistance to inhibition by EBV BNLF2a. Similar to Figure 3A, histograms indicate the abundance of cells with different intensity of anti-MHC-I cell surface stain in cells expressing BNLF2a with the indicated variants.

## DISCUSSION

ICP47 binds to TAP and blocks its function, enabling HSV1 infected cells to evade the host immune system [7,14]. In the present study, we tested a library of variants in the TAP2 subunit of the TAP peptide transporter for resistance to the HSV1 ICP47 protein. The effects of the ICP47 inhibition were scored by measuring the abundance of MHC-I on the cell surface. Out of the 1786 variants present in the multiplexed library, 39 were ICP47-resistant.

Another study applied an analogous approach to rescue TAP2 KO cells with eight variants in TAP2 detected in the human population [15]. In that study, all eight of the variants functioned similarly in TAP transport, and they all were similarly sensitive to inhibition by ICP47, US6, and BNLF2a. By contrast, in the current study we used a multiplexed approach to generate 1786 TAP2 variants, which were functionally screened for resistance to ICP47 in regulating MHC-I cell surface expression. In our multiplexed screen, we captured 811 variants, and 39 were found to be resistant to ICP47.

Some of the TAP2 substitutions giving rise to ICP47-resistant variants were at residues (Ser274, Ser254, Thr244 and Thr257) that mapped near the ICP47 binding site. Other substitutions (Ser289 and Thr293) are distant to the binding site, and others (Thr511 and Ser635) map in the TAP2 NTP binding domain. Since the original TAP2 variant library targeted 96 codons throughout the length of the 686 amino acid TAP2 protein, and since most of the substitutions giving rise to ICP47-resistant variants mapped on the 3D structure near the ICP47 binding site, we infer that some of these substitutions may affect the binding of ICP47 to the TAP complex. The two substitutions (T293P and S289L) that are distant to the ICP47 binding site but give rise to ICP47-resistant variants, may also affect the TAP/ICP47 interaction by inducing conformational changes altering the architecture of the interface between the two interacting partners. The same may also apply to the substitutions that map in the TAP2 NTP-binding domain.

Different Herpes virus family members use different proteins to bind and inactivate the TAP peptide transporter, raising the question whether their immunosuppressive functions mechanistically overlap. The ICP47-resistant variants of TAP2 gave us the opportunity to directly address this question. Expression of the EBV BNLF2a protein in TAP2 knockout U2OS cells rescued with wild type TAP2, or four different ICP47-resistant TAP2 variants, revealed that the ICP47-resistant TAP2 variants and the wild type TAP2 were equally sensitive to inhibition by BNLF2a. These results provided unequivocal evidence that although both ICP47 and BNLF2a inhibit TAP by binding the TAP complex, the precise mechanisms of the inhibitory function of the HSV1 and EBV proteins differ. It would be interesting to repeat the multiplexed library selection for TAP2 variants resistant to BNLF2a to compare BNLF2a-resistant variants with those resistant to ICP47.

The 3D structures have been determined for TAP bound to ICP47 [9,10] and TAP bound to the peptide cargo (PDB 8T4J) (Figure 5). Based on these structures, the ICP47 protein makes extensive contacts with TAP along its inward facing channel of TAP, while the antigenic peptide binds to TAP via its ends, which contact a small set of residues in the inward facing channel of TAP. Some substitutions giving rise to ICP47-resistant variants impact strongly the ICP47 binding site (Figure 5, *left*) but only one (S274H) maps near the peptide binding site (Figure 5, *right*), and that one should not affect peptide binding because the variant carrying this substitution is functionally normal. The fact that we identified substitutions which block ICP47 sensitivity, without affecting TAP peptide transport, suggests that it would be feasible to isolate small molecule inhibitors of ICP47, that will not affect the normal function of TAP. Such molecules should be therapeutically effective as antiviral agents. Based on the structures in Figure 5, a compound that binds to TAP2 residues 244-257 could impair ICP47 binding but not peptide binding and could thus be the basis of a new antiviral strategy for herpes virus infection.

**Figure 5.**
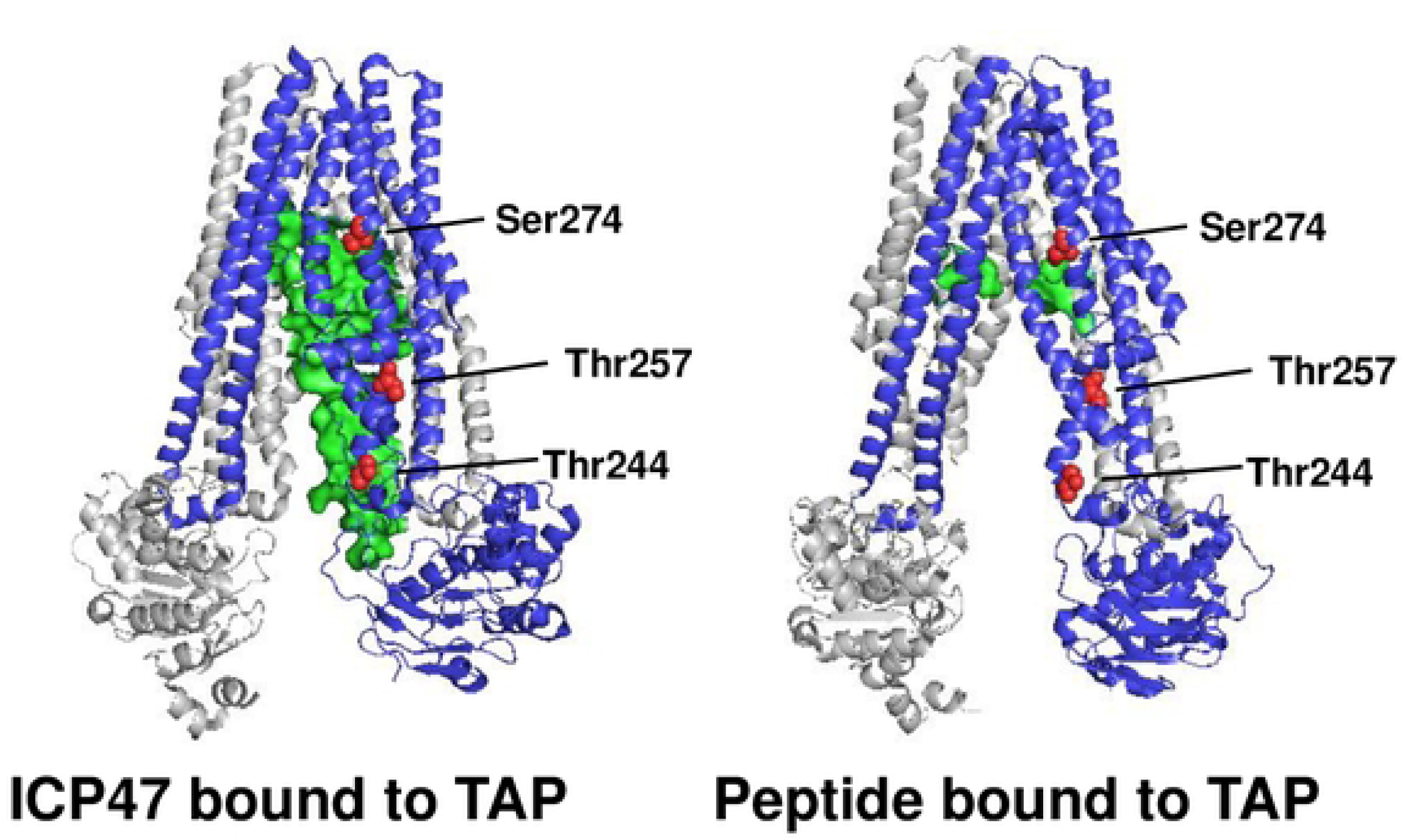
Comparison of 3D structures of TAP bound to ICP47 and TAP bound to antigenic peptide. The TAP transporter in bound to ICP47 (PDB 5U1D) [9,10] or bound to peptide cargo (PDB 8T4J) are shown with three TAP2 residues at which variants were resistant to ICP47 inhibition are shown in red. Image drawn with PyMOL.

## MATERIALS AND METHODS

### Plasmids and cell lines

All experiments utilized cell lines derived from U2OS (ATCC catalog HTB-22) that had integrated at a single locus in the genome an attP sequence downstream of the RSV-LTR [12]. The TAP2 gene was knocked out by CRISPR and knock out cells were selected by labeling cells with anti-MHC-I (see below) and collecting cells that had low surface stain of MHC-I by flow cytometry using the Cytek Aurora instrument in the Ohio State University Comprehensive Cancer Center Flow Cytometry shared resource. TAP2-knock out cells were rescued for MHC-I surface expression by transfection of a pUC-19 plasmid backbone that contained an attB sequence upstream of the hygromycin resistance gene, the P2A translation re-start sequence, and the TAP2 cDNA. Since this plasmid, pUC19-attB-Hygro-TAP2, lacked a promoter for mammalian cells, hygromycin resistance and TAP2 were only expressed in cells in which the integrated attP site recombined with the plasmid encoded attB site (manuscript submitted [11]). A PCR-based approach [16] was employed to target 96 codons of TAP2 using degenerate oligonucleotides that replaced the wild-type codon with NNK sequence (N= A, T, G, or C; K = G or T) and an 18 bp barcode in the 3’-UTR. Each plasmid molecule contained wild-type sequence everywhere except at the targeted codon, and each variant molecule had a barcode that could identify the variant. The plasmid library was confirmed by long read sequencing with a PacBio instrument (Drexel University sequencing facility) to have 1786 variants, each variant with a specific barcode [11]. Transfection into the U2OS TAP2 knock-out cells of this library, growth in the presence of doxycycline (10 µg/ml) and selection for hygromycin resistance (50 µg/ml) resulted in a pool of cells that integrated the plasmid library into the genome, one variant per cell.

For expression of ICP47 or BNLF2a, pHAGE2-EF1aL-dsRed-W (Addgene plasmid #195863, a gift from Darrell Kotton) was modified to remove the stop codon on the dsRed coding sequence and insert either P2A-ICP47 or P2A-BNLF2a DNA sequences using a gene block (IDT). The gene block was inserted into the pHAGE2 vector using the NEBuilder HiFi DNA Assembly Cloning kit. Sequences of plasmid products were confirmed using the Ohio State University Comprehensive Cancer Center Genomics shared resource. Lentivirus was prepared using standard procedures by co-transfecting into HEK293T cells pHAGE2-EF1aL-dsRed-ICP47 or pHAGE2-EF1aL-dsRed-BNLF2a along with plasmids for expressing lentiviral structural proteins; virus was collected in the supernatant medium. Virus titers were determined by counting with flow cytometry, using the BD Fortessa instrument, the dsRed-positive cells when the virus was infected into the U2OS cell line.

For generation of selected TAP2 variants for singleton analysis, oligonucleotides encoding the targeted sequence change (IDT) were subcloned into the pUC13-attB-Hygro-TAP2 plasmid using the HiFi kit (New England Biolabs). After confirmation of the sequence change, the plasmid was transfected into the U2OS TAP2-knock-out cells and recombined into the integrated attP site as described above; and these cells were selected by growth in 50 µg/ml hygromycin and 10 µg/ml doxycycline.

### Selection of TAP2 variants resistant to ICP47

Cells containing the integrated library of TAP2 variants were infected at an MOI of 0.5 with the titered ICP47-expressing lentivirus, and five days post infection cells were stained with Brilliant Ultra Violet 395-conjugated anti-HLA A/B/C antibody (anti-MHC-I, Thermo-Fisher) and analyzed by flow cytometry using the Cytek Aurora instrument. Cells that were dsRed-positive and expressing high concentration of MHC-I per cell were collected. These cells (sort 1) were grown in tissue culture to amplify the cell number and subjected to two more rounds of sorting for high MHC-I expressers and re-amplification. After sort 3, the genomic DNA was isolated from the collected cells, the barcode was amplified by two rounds of PCR using established methods [17,18], and the sample was analyzed by sequencing using the Illumina instrument at the Memorial Sloan-Kettering Cancer Center sequencing facility.

Cells expressing targeted single mutations of TAP2 were infected with the ICP47-expressing or BNLF2a-expressing lentivirus at an MOI of 0.5 and analyzed five days post-infection by staining with the Brilliant Ultra Violet 395-conjugated anti-HLA A/B/C antibody as described above and analyzed by flow cytometry using the BD Fortessa instrument in the Ohio State University Comprehensive Cancer Center Flow Cytometry shared resource.

### Statistical analysis of TAP2 variants enriched in the ICP47-resistant pool of cells

We conducted a statistical analysis using a multinomial distribution to assess whether specific variants occurred more frequently than expected by chance in the barcode sequencing data. Barcodes for each variant were decoded with Enrich2 [19], and variants with missing values were excluded, yielding a dataset of 811 variants. The total sequencing depth was calculated as the sum of counts across all variants. Under the null hypothesis that each variant had an equal probability of occurrence, we ran 100,000 simulations to model the expected distribution of variant counts. The p-value for each variant was calculated as the proportion of simulations where the randomly allocated count met or exceeded the observed count.

In our statistical inference, assuming equal probability for all variants is a conservative approach because, in reality, some variants naturally occur more frequently due to biological factors. Ignoring these differences may assign higher p-values to truly enriched variants, potentially leading to false negatives. However, this conservative approach ensures that only the most definitively enriched variants stand out with low p-values, resulting in binary p-values of 0 or 1. Therefore, in the inference, a p-value of 0 indicates significant enrichment beyond random chance, suggesting potential biological significance.

## ACKNOWLEDGMENTS

This work was supported by NIH R01 CA228083 to JDP. GN was supported by a Pelotonia Graduate Training Fellowship. Research reported in this publication was supported by The Ohio State University Comprehensive Cancer Center and the National Institutes of Health under grant number P30 CA016058.

## LEGENDS TO SUPPLEMENTARY FIGURES

**Figure S1. Scatterplots of flow cytometry results from U2OS cells expressing TAP2 variants for effects of ICP47 on MHC-I surface abundance.**

In each scatterplot, the x-axis is the intensity of MHC-I antibody stain, and the y-axis is the intensity of dsRed stain. Uninfected cells are shown on the left, and cells expressing dsRed and ICP47 are shown on the right. Each row is a different variant of TAP2: TAP2-wild-type, TAP2-S274H, TAP2-Y537P, TAP2-T257I, TAP2-T244R, and TAP2-T499D, as indicated. Each cell line is a U2OS-derived cell with TAP2 knocked out by CRISPR and rescued by the stable integration of the indicated variant at a single recombination site in the genome.

**Figure S2. Scatterplots of flow cytometry results from U2OS cells expressing TAP2 variants for effects of BNLF2a on MHC-I surface abundance.**

As in Figure S1, each scatterplot measures the intensity of MHC-I antibody stain (x-axis) and dsRed (y-axis). The left column contains results from ICP47-expressing cells (a different experiment than used in Figure S1) and the right column contains results from the same cells but expressing BNLF2a.

**Figure S3. Full images of immunoblots**

Full exposures are shown for immunoblots shown in Figure 3C.

## LEGENDS TO SUPPLEMENTARY TABLES

**Table S1. Read-count frequency for each TAP2 variant in cells with high MHC-I surface stain.**

TAP2 variants detected in the high MHC-I surface stain cells are listed in order of the number of read-counts for each variant. The six highest are indicated in red, the next six in yellow, and the top 39 are indicated with color highlights. The same colors are used to label the residues in the 3D structure in Figure 2C. Synonymous and nonsense variants are highlighted in gray.

**Table S2. Functional scores of variants independent of a viral inhibitor protein.**

The functional scores of 1405 variants were scored in prior experiments [11]. Functional scores less than 0.92 were judged loss of function, and functional scores greater than 2.2 were judged hyperfunctional. The 39 variants that were significantly enriched in the pool of TAP2 variants are shown with their functional scores and interpretation.

**Table S3. Quantitation of singleton assays for TAP2 variant resistance to inhibition by ICP47 or BNLF2a**

Results of the median scores for variant effects on MHC-I surface abundance are indicated for each replicate. The ratio of the median in the presence of ICP47 to the median in the absence of ICP47 was used to compare variants to wild-type and was used in the table in Figure 3B.

## Notes

### Competing Interest Statement

The authors have declared no competing interest.

## REFERENCES

[1] Dhatchinamoorthy, K., Colbert, J.D. and Rock, K.L. (2021) Cancer Immune Evasion Through Loss of MHC Class I Antigen Presentation. Frontiers in Immunology, 12. 10.3389/fimmu.2021.636568

[2] Mantel, I., Sadiq, B.A. and Blander, J.M. (2022) Spotlight on TAP and its vital role in antigen presentation and cross-presentation. *Molecular Immunology*, Elsevier Ltd. 142, 105–19. 10.1016/j.molimm.2021.12.013

[3] Parcej, D. and Tampé, R. (2010) ABC proteins in antigen translocation and viral inhibition. Nature Chemical Biology, 6, 572–80. 10.1038/nchembio.410

[4] Eggensperger, S. and Tampé, R. (2015) The transporter associated with antigen processing: A key player in adaptive immunity. Biological Chemistry, 396, 1059–72. 10.1515/hsz-2014-0320

[5] Gadola, S.D., Moins-Teisserenc, H.T., Trowsdale, J., Gross, W.L. and Cerundolo, V. (2000) TAP deficiency syndrome. Clin. Exp. Immunol. 10.1046/j.1365-2249.2000.01264.x

[6] DeSandro, A.M., Nagarajan, U.M. and Boss, J.M. (2000) Associations and Interactions between Bare Lymphocyte Syndrome Factors. Molecular and Cellular Biology, 20, 6587 LP – 6599. 10.1128/MCB.20.17.6587-6599.2000

[7] Neumann, L., Kraas, W., Uebel, S., Jung, G. and Tampé, R. (1997) The active domain of the herpes simplex virus protein ICP47: A potent inhibitor of the transporter associated with antigen processing (TAP). Journal of Molecular Biology, 272, 484–92. 10.1006/jmbi.1997.1282

[8] Verweij, M.C., Horst, D., Griffin, B.D., Luteijn, R.D., Davison, A.J., Ressing, M.E. and Wiertz, E.J.H.J. (2015) Viral Inhibition of the Transporter Associated with Antigen Processing (TAP): A Striking Example of Functional Convergent Evolution. PLoS Pathogens, 11, 1–19. 10.1371/journal.ppat.1004743

[9] Oldham, M.L., Grigorieff, N. and Chen, J. (2016) Structure of the transporter associated with antigen processing trapped by herpes simplex virus. ELife, 5, 1–16. 10.7554/eLife.21829

[10] Oldham, M.L., Hite, R.K., Steffen, A.M., Damko, E., Li, Z., Walz, T. and Chen, J. (2016) A mechanism of viral immune evasion revealed by cryo-EM analysis of the TAP transporter. Nature, Nature Publishing Group. 529, 537–40. 10.1038/nature16506

[11] Nagy, G., Presley, R., Jeon, H., Moreira, L., Dhar, S., Tull, O., Soysal, B., Yadati, A.S., Cook, K.D., Schnell, L.E., Gupta, A., Banerjee, T., Chung, D., Tsichlis, P.N. and Parvin, J.D. (2024) Multiplexed functional analysis of TAP2 variants in regulating MHC-I cell surface abundance reveals overexpression of PLK1 downregulates antigen presentation. BioRxiv, 2024.09.30.615919. 10.1101/2024.09.30.615919

[12] Matreyek, K.A., Stephany, J.J., Chiasson, M.A., Hasle, N. and Fowler, D.M. (2020) An improved platform for functional assessment of large protein libraries in mammalian cells. Nucleic Acids Research, Oxford University Press. 48, 1–12. 10.1093/nar/gkz910

[13] Horst, D., van Leeuwen, D., Croft, N.P., Garstka, M.A., Hislop, A.D., Kremmer, E., Rickinson, A.B., Wiertz, E.J.H.J. and Ressing, M.E. (2009) Specific Targeting of the EBV Lytic Phase Protein BNLF2a to the Transporter Associated with Antigen Processing Results in Impairment of HLA Class I-Restricted Antigen Presentation. The Journal of Immunology, 182, 2313–24. 10.4049/jimmunol.0803218

[14] Galocha, B.B., Hill, A., Barnett, B.C., Dolan, A., Raimondi, A., Cook, R.F., Brunner, J., Mcgeoch, D.J. and Ploegh, H.L. (1997) Inhibitor of the Major Histocompatibility Complex. Peptides, 185, 1565–72.

[15] Praest, P., Luteijn, R.D., Brak-Boer, I.G.J., Lanfermeijer, J., Hoelen, H., Ijgosse, L., Costa, A.I., Gorham, R.D., Lebbink, R.J. and Wiertz, E.J.H.J. (2018) The influence of TAP1 and TAP2 gene polymorphisms on TAP function and its inhibition by viral immune evasion proteins. *Molecular Immunology*, Elsevier. 101, 55–64. 10.1016/j.molimm.2018.05.025

[16] Jain, P.C. and Varadarajan, R. (2014) A rapid, efficient, and economical inverse polymerase chain reaction-based method for generating a site saturation mutant library. Analytical Biochemistry, 449, 90–8. 10.1016/j.ab.2013.12.002

[17] Adamovich, A.I., Diabate, M., Banerjee, T., Nagy, G., Smith, N., Duncan, K., Mendoza Mendoza, E., Prida, G., Freitas, M.A., Starita, L.M. and Parvin, J.D. (2022) The functional impact of BRCA1 BRCT domain variants using multiplexed DNA double-strand break repair assays. *American Journal of Human Genetics*, American Society of Human Genetics. 109, 618–30. 10.1016/j.ajhg.2022.01.019

[18] Nagy, G., Diabate, M., Banerjee, T., Adamovich, A.I., Smith, N., Jeon, H., Dhar, S., Liu, W., Burgess, K., Chung, D., Starita, L.M. and Parvin, J.D. (2023) Multiplexed assay of variant effect reveals residues of functional importance in the BRCA1 coiled-coil and serine cluster domains. PLoS ONE, 18, e0293422. 10.1371/journal.pone.0293422

[19] Rubin, A.F., Gelman, H., Lucas, N., Bajjalieh, S.M., Papenfuss, A.T., Speed, T.P. and Fowler, D.M. (2017) A statistical framework for analyzing deep mutational scanning data. Genome Biology, 18. 10.1186/s13059-017-1272-5

